# Improved Neutralization of Omicron BA.4/5, BA.4.6, BA.2.75.2, BQ.1.1, and XBB.1 with Bivalent BA.4/5 Vaccine

**DOI:** 10.1101/2022.11.17.516898

**Authors:** Jing Zou, Chaitanya Kurhade, Sohil Patel, Nicholas Kitchin, Kristin Tompkins, Mark Cutler, David Cooper, Qi Yang, Hui Cai, Alexander Muik, Ying Zhang, Dung-Yang Lee, Ugur Sahin, Annaliesa S. Anderson, William C. Gruber, Xuping Xie, Kena A. Swanson, Pei-Yong Shi

**Author notes:** These authors contribute equally to this study (Jing Zou, Chaitanya Kurhade). Corresponding authors: X.X., K.A.S., P.-Y.S.

## Abstract

The BNT162b2 bivalent BA.4/5 COVID-19 vaccine has been authorized to mitigate COVID-19 due to current Omicron and potentially future variants. New sublineages of SARS-CoV-2 Omicron continue to emerge and have acquired additional mutations, particularly in the spike protein, that may lead to improved viral fitness and immune evasion. The present study characterized neutralization activities against new Omicron sublineages BA.4.6, BA.2.75.2, BQ.1.1, and XBB.1 after a 4^th^ dose (following three doses of BNT162b2) of either the original monovalent BNT162b2 or the bivalent BA.4/5 booster in individuals >55 years of age. For all participants, the 4^th^ dose of monovalent BNT162b2 vaccine induced a 3.0×, 2.9×, 2.3×, 2.1×, 1.8×, and 1.5× geometric mean neutralizing titer fold rise (GMFR) against USA/WA1-2020 (a strain isolated in January 2020), BA.4/5, BA.4.6, BA.2.75.2, BQ.1.1, and XBB.1, respectively; the bivalent vaccine induced 5.8×, 13.0×, 11.1×, 6.7×, 8.7×, and 4.8× GMFRs. For individuals without SARS-CoV-2 infection history, BNT162b2 monovalent induced 4.4×, 3.0×, 2.5×, 2.0×, 1.5×, and 1.3× GMFRs, respectively; the bivalent vaccine induced 9.9×, 26.4×, 22.2×, 8.4×, 12.6×, and 4.7× GMFRs. These data suggest the bivalent BA.4/5 vaccine is more immunogenic than the original BNT162b2 monovalent vaccine against circulating Omicron sublineages, including BQ.1.1 that is becoming prevalent globally.

## Main text

Severe acute respiratory syndrome coronavirus 2 (SARS-CoV-2) Omicron variant continues to evolve globally into sublineages since its emergence in November 2021^1^. To mitigate the ongoing Omicron pandemic, the U.S. FDA and European Medicines Agency authorized emergency use of the BNT162b2 bivalent BA.4/5-vaccine (BA.4 and BA.5 encode an identical spike protein sequence) in September 2022; the vaccine was subsequently authorized for use in many countries globally. The bivalent vaccine contains two mRNAs: one encoding the wild-type Wuhan SARS-CoV-2 spike protein included in the BNT162b2 monovalent vaccine (or original vaccine) and another encoding the Omicron sublineage BA.4/5-spike protein. Currently, new Omicron BA.2- and BA.4/BA.5-descendent sublineages (e.g., BA.4.6, BA.2.75.2, BQ.1.1, and XBB.1) have emerged (https://covid.cdc.gov/covid-data-tracker/#variant-proportions). Although early epidemiological data suggest these new sublineages have not shown signs of increased disease severity, they have accumulated additional spike mutations that could further evade vaccine- and/or infection-elicited antibody neutralization^2–4^. Here we compared the neutralization activities against these Omicron sublineages after a 4^th^ dose (following three doses of BNT162b2) of either the original monovalent BNT162b2 or the bivalent BA.4/5 booster in humans.

Participants >55-years-old received 3 prior doses of 30-μg BNT162b2 and subsequently a 4^th^ dose booster of 30-μg monovalent BNT162b2 at ~6.6-months-post-dose-3 (Study C4591031) or bivalent vaccine (15-μg BNT162b2 plus 15-μg BA.4/5) at ~11-months-post-dose-3 (Study C4591044). Serum was collected on the day of dose 4 (Pre serum) and at 1-month-post-dose-4 (1MPD4 serum). All participants were tested for evidence of SARS-CoV-2 infection by viral nucleocapsid antibodies and RT-PCR test; a subset of participants from both vaccine groups, equally distributed among those with or without evidence of infection at baseline (Pre serum), were selected for the neutralization analysis. For neutralization testing, the complete spike gene from Omicron BA.4/5, BA.4.6, BA.2.75.2, BQ.1.1, or XBB.1 was engineered into the backbone of mNeonGreen (mNG) reporter USA-WA1/2020 SARS-CoV-2 (a strain isolated in January 2020)^5^. The resulting wild-type- (WT), BA.4/5-, BA.4.6, BA.2.75.2-, BQ.1.1-, and XBB.1-spike mNG USA-WA1/2020 were used to measure 50% fluorescent focus reduction neutralization titers (FFRNT_50_) for each serum. **Tables S1** and **S2** summarize the serum information and their FFRNT_50_ values, respectively.

For all participants, the 4^th^ dose of monovalent BNT162b2 vaccine induced a 3.0×, 2.9×, 2.3×, 2.1×, 1.8×, and 1.5× geometric mean neutralizing titer fold rise (GMFR) against WT, BA.4/5, BA.4.6, BA.2.75.2, BQ.1.1, and XBB.1, respectively; the bivalent vaccine induced 5.8×, 13.0×, 11.1×, 6.7×, 8.7×, and 4.8× GMFRs (**Fig. 1A**). For individuals without SARS-CoV-2 infection history, BNT162b2 monovalent induced 4.4×, 3.0×, 2.5×, 2.0×, 1.5×, and 1.3× GMFRs, respectively; the bivalent vaccine induced 9.9×, 26.4×, 22.2×, 8.4×, 12.6×, and 4.7× GMFRs (**Fig. 1B**). For individuals with previous SARS-CoV-2 infection, BNT162b2 monovalent induced 2.0×, 2.8×, 2.1×, 2.1×, 2.2×, and 1.8× GMFRs, respectively; the bivalent vaccine induced 3.5×, 6.7×, 5.6×, 5.3×, 6.0×, and 4.9× GMFRs (**Fig. 1C**). Despite different intervals from dose 3 to 4, the pre-dose-4 neutralizing titers were similar between the monovalent and bivalent vaccine groups in all the participants, regardless of SARS-CoV-2 infection history.

**Figure 1.**
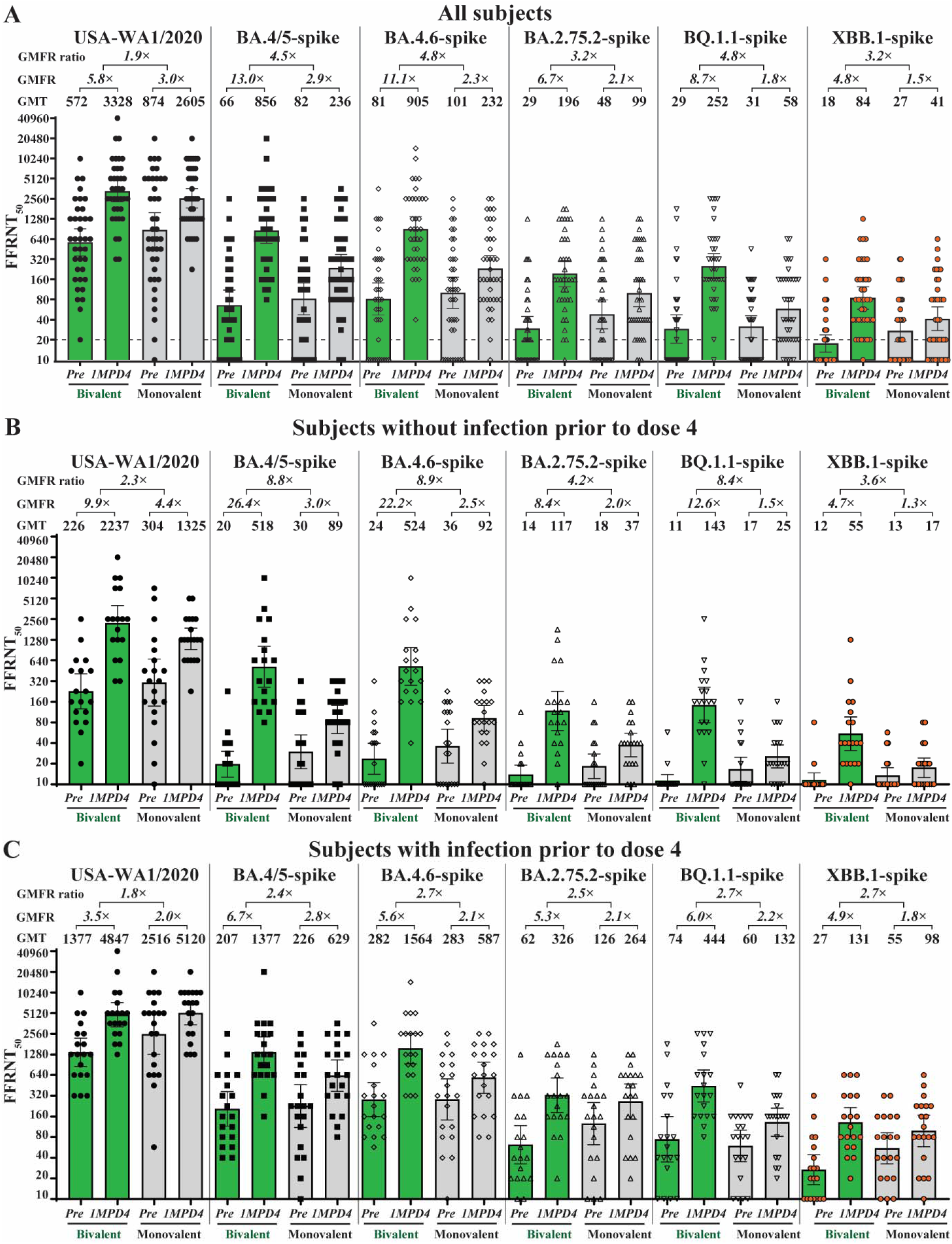
USA-WA1/2020, Omicron BA.4/5, BA.4.6, BA.2.75.2, BQ.1.1, and XBB.1 neutralizing response with bivalent BA.4/5 or monovalent BNT162b2 booster. The bar heights and the numbers above indicate geometric means of neutralization titers (GMTs). The whiskers indicate 95% CI. Green bars, 4^th^ dose vaccination with bivalent; gray bars, 4^th^ dose vaccination with BNT16b2 monovalent. FFRNT_50_s against USA-WA1/2020, BA.4/5-spike, BA.4.6-spike, BA.2.75.2-spike, BQ.1.1-spike, and XBB.1-spike are shown as black circle, black square, rhombus, up-triangle, down-triangle, and red circle with black outline, respectively. Pre, serum samples collected on the day of booster; 1MPD4, one month post dose 4. GMFR, geometric mean fold rises (ratio of titers at 1MPD4 relative to Pre). Numbers above GMFRs indicate the ratios between GMFRs of bivalent to GMFRs of monovalent. (A) FFRNT_50_s of all subjects regardless of infection status. (B) FFRNT_50_s of all subjects without evidence of SARS-CoV-2 infection prior to dose 4. (C) FFRNT_50_s of all subjects with evidence of SARS-CoV-2 infection prior to dose 4. For panels A, B, and C, the lower bounds of the two-sided 95% CIs for GMFR of bivalent or monovalent booster against USA-WA1/2020, BA.4/5-spike, BA.4.6, BA.2.75.2, BQ.1.1, and XBB.1 were all >1. The two-sided 95% CIs for GMFR of bivalent and monovalent booster against USA-WA1/2020, BA.4/5-spike, BA.4.6, BA.2.75.2, BQ.1.1, and XBB.1 were all nonoverlapping (Table S4).

Our results support three conclusions. First, the bivalent BA.4/5 vaccine consistently elicited higher neutralizing responses against BA.5-derived sublineages (BA.4.6, BQ.1.1, and XBB.1) and BA.2-derived sublineage (BA.2.75.2) than the original BNT162b2 monovalent vaccine when administered as a 4^th^ dose booster, regardless of SARS-CoV-2 infection history. Second, individuals with SARS-CoV-2 infection history developed higher neutralizing titers than those without infection history after the 4^th^ dose booster. Third, for each tested Omicron sublineage, the difference between the monovalent original and bivalent GMFR was greater for the group of sera without previous infection than the group with previous infection.

Among all Omicron sublineages, BA.2.75.2, BQ.1.1, and XBB.1 exhibit the lowest vaccine-elicited neutralization; however, neutralizing titers following a bivalent booster were several fold higher than those following the original BNT162b2 vaccine. These data suggest the bivalent vaccine is more immunogenic than the original vaccine against circulating Omicron sublineages, supporting current bivalent vaccine use, and underscore the importance of monitoring real-world effectiveness.

## Acknowledgments

We thank the Pfizer-BioNTech clinical trial C4591031 and C4591044 participants, from whom the sera were obtained. We thank the many colleagues at Pfizer and BioNTech who developed and produced the BNT162b2 and bivalent vaccine.

## Author contributions

Conceptualization, D.C., X.X., K.A.S. P.-Y.S.; Methodology, J.Z., C.K., Q.Y., X.X., P.-Y.S.; Investigation, J.Z., C.K., Q.Y., M.C., D.C., A.M., S.P., A.S.A., N.K., X.X., K.A.S., P.-Y.S.; Resources, M.C., D.C., A.M., U.S., X.X., A.S.A.,K.A.S., P.-Y.S.; Data Curation, J.Z., C.K., Y.Z., X.X., K.T.; Writing-Original Draft, X.X., K.A.S., P.-Y.S.; Writing-Review & Editing, J.Z., C.K., Q.Y., H.C., M.C., D.C., A.S.A., W.C.G., N.K., A.M., U.S., X.X., K.A.S., P.-Y.S, Y.Z., D.-Y.L., K.T., S.P.; Supervision, A.S.A., W.C.G., N.K, K.A.S., X.X., P.-Y.S.; Funding Acquisition, A.S.A., D.C., X.X., K.A.S., P.-Y.S.

