## Supplementary table 1-4 for "Improved Neutralization of Omicron BA.4/5, BA.4.6, BA.2.75.2, BQ.1.1, and XBB.1 with Bivalent BA.4/5 Vaccine"

**Supplementary materials**

### Methods

#### Biosafety level-3 operation

All virus work was performed in a biosafety level 3 (BSL-3) laboratory with redundant fans in the biosafety cabinets at the University of Texas Medical Branch at Galveston. All personnel wore powered air-purifying respirators (Breathe Easy, 3M) with Tyvek suits, aprons, booties, and double gloves.

#### Cells

Vero E6 (ATCC® CRL-1586) purchased from the American Type Culture Collection (ATCC, Bethesda, MD) and Vero E6 cells expressing TMPRSS2 purchased from SEKISUI XenoTech, LLC were maintained in a high-glucose Dulbecco's modified Eagle's medium (DMEM) containing 10% fetal bovine serum (FBS; HyClone Laboratories, South Logan, UT) and 1% penicillin/streptomycin at 37°C with 5% $CO_2$. Culture media and antibiotics were purchased from Thermo Fisher Scientific (Waltham, MA). The cell line was tested negative for *Mycoplasma*.

#### Human Serum

In participants >55 years of age who previously received three 30-µg BNT162b2 doses, serum samples were collected just prior to and 1 month post-boost with a 4$^{th}$ dose booster of monovalent original 30-µg BNT162b2 (Study C4591031; ClinicalTrials.gov identifier: NCT04955626) or 30-µg BNT162b2 bivalent BA.4/BA.5 vaccine (15 µg original with 15 µg BA.4/BA.5) (Study C4591044; ClinicalTrials.gov identifier: NCT05472038). The protocol and informed consent were approved by institutional review boards for each of the investigational centers participating in the study. The study was conducted in compliance with all International Council for Harmonisation Good Clinical Practice guidelines and the ethical principles of the Declaration of Helsinki. . The median time between dose-3 and -4 for Study C4591031 and C4591044 was 6.3 and 11.3 months, respectively. All participants were screened by SARS-CoV-2 nucleocapsid Ig serological test for preexisting SARS-CoV-2 or RT-PCR for existing infection. The subset of participant's sera (approximately 40 per vaccine group) for neutralization testing were selected to ensure equal distribution between those with and without evidence of infection at baseline (pre-dose-4) by either test and medical history of COVID-19. **Table S1** summarizes the detailed demographic information (e.g., age and gender) of all participants. Human sera were heat-inactivated at 56°C for 30 min before the neutralization test.

**Recombinant Omicron sublineages-mNG SARS CoV-2 viruses**

Recombinant Omicron sublineage BA.4/5-, BA.4.6-, BA.2.75.2-, BQ.1.1-, and XBB.1-spike mNG SARS-CoV-2s was constructed by engineering the complete spike gene from the indicated variants into an infectious cDNA clone of mNG USA-WA1/2020 and reported previously[1-5]. Viruses were rescued post 2-3 days after electroporation and served as P0 stock. P0 stock was further passaged once on Vero E6 cells to produce P1 stock. The spike gene was sequenced from all P1 stock viruses to ensure no undesired mutation. The infectious titer of the P1 virus was quantified by fluorescent focus assay on Vero E6 cells. The P1 virus was used for the neutralization test.

**Fluorescent focus reduction neutralization test (FFRNT)**

Neutralization titers of human sera were measured by FFRNT using the USA-WA1/2020-, BA.4/5, BA.4.6-, BA.2.75.2-, BQ.1.1- and XBB.1-spike mNG SARS-CoV-2s. All sera were tested sequentially, USA-WA1/2020 and BA.4/5 followed by the remaining Omicron sublineages. The details of the FFRNT protocol were reported previously[1,5-8]. Briefly, $2.5 \times 10^4$ Vero E6 cells per well were seeded in 96-well plates (Greiner Bio-one™). The cells were incubated overnight. On the next day, each serum was 2-fold serially diluted in the culture medium with the first dilution of 1:20 (final dilution range of 1:20 to 1:20,480). The diluted serum was incubated with 100-150

FFUs of mNG SARS-CoV-2 at 37 °C for 1 h, after which the serum virus mixtures were loaded onto the pre-seeded Vero E6 cell monolayer in 96-well plates. After 1 h infection, the inoculum was removed and 100 µl of overlay medium (supplemented with 0.8% methylcellulose) was added to each well. After incubating the plates at 37 °C for 16 h, raw images of mNG foci were acquired using Cytation™ 7 (BioTek) armed with 2.5× FL Zeiss objective with a wide-field of view and processed using the Gene 5 software settings (GFP [469,525] threshold 4000, object selection size 50-1000 µm). The foci in each well were counted and normalized to the non-serum-treated controls to calculate the relative infectivities. The $FFRNT_{50}$ value was defined as the minimal serum dilution that suppressed >50% of fluorescent foci. The neutralization titer of each serum was determined in duplicate assays, and the geometric mean was taken. All attempts at replication were successful. Supplementary Table 2 summarizes the $FFRNT_{50}$ results. Data were initially plotted in GraphPad Prism 9 software and assembled in Adobe Illustrator.

Supplementary Table 1. Demographics information of subjects.

| Characteristic | Vaccine Group: C4591044 BNT162b2 Bivalent (WT/OMI BA.4/BA.5) 30 µg (N[a]=38) n[b] (%) | Vaccine Group: C4591031 BNT162b2 30 µg (N[a]=40) n[b] (%) |
|---|---|---|
| Sex | | |
| Male | 24 (63.2) | 17 (42.5) |
| Female | 14 (36.8) | 23 (57.5) |
| Age at vaccination (years) | | |
| Mean (SD) | 66.5 (6.82) | 65.4 (5.80) |
| Median | 66.0 | 65.5 |
| Min, max | (57, 79) | (56, 79) |
| Baseline SARS-CoV-2 status | | |
| Positive[c] | 19 (50.0) | 20 (50.0) |
| Negative[d] | 19 (50.0) | 20 (50.0) |
| Time from the last dose of BNT162b2 (received prior to the study) to the study vaccination (months) | | |
| n | 38 | 40 |
| Mean (SD) | 10.9 (1.41) | 6.9 (1.82) |
| Median | 11.3 | 6.3 |
| Min, max | (5.5, 13.0) | (5.4, 13.1) |
| ≥5 to <7 Months | 1 (2.6) | 30 (75.0) |
| ≥7 to <9 Months | 2 (5.3) | 5 (12.5) |
| ≥9 to ≤12 Months | 31 (81.6) | 4 (10.0) |
| >12 Months | 4 (10.5) | 1 (2.5) |

Abbreviations: N-binding = SARS-CoV-2 nucleoprotein–binding; NAAT = nucleic acid amplification test

a. N = number of participants in the specified group, or the total sample. This value is the denominator for the percentage calculations.
b. n = Number of participants with the specified characteristic.
c. Positive N-binding antibody result at baseline, positive NAAT result at baseline, or medical history of COVID-19.
d. Negative N-binding antibody result at baseline, negative NAAT result at baseline, and no medical history of COVID-19.

Supplementary Table 2. FFRNT$_{50}$ values of all subjects received bivalent BA.4/5 booster.

| ID | AGE (years) | SEX | Booster | *FFRNT$_{50}$ | | | | | | | | | | | | With infection at baseline (pre Dose 4) |
|---|---|---|---|---|---|---|---|---|---|---|---|---|---|---|---|---|
| | | | | USA-WA1/2020 | | BA.4/5-spike | | BA.4.6-spike | | BA.2.75.2-spike | | BQ.1.1-spike | | XBB.1-spike | | |
| | | | | Pre | 1MPD4 | Pre | 1MPD4 | Pre | 1MPD4 | Pre | 1MPD4 | Pre | 1MPD4 | Pre | 1MPD4 | |
| 1 | 73 | M | Bivalent | 5120 | 5120 | 640 | 2560 | 1280 | 5120 | 320 | 905 | 320 | 1280 | 57 | 160 | YES |
| 2 | 78 | M | Bivalent | 640 | 20480 | 80 | 3620 | 160 | 5120 | 40 | 1280 | 40 | 1810 | 20 | 640 | YES |
| 3 | 76 | M | Bivalent | 1280 | 7241 | 40 | 905 | 80 | 640 | 40 | 160 | 10 | 320 | 10 | 80 | NO |
| 4 | 61 | F | Bivalent | 905 | 3620 | 113 | 640 | 113 | 320 | 10 | 20 | 57 | 113 | 10 | 20 | YES |
| 5 | 71 | M | Bivalent | 113 | 640 | 20 | 160 | 10 | 160 | 10 | 40 | 10 | 57 | 10 | 20 | NO |
| 6 | 68 | M | Bivalent | [$]N/A | N/A | N/A | N/A | N/A | N/A | N/A | N/A | N/A | N/A | N/A | N/A | NA |
| 7 | 57 | F | Bivalent | 3620 | 3620 | 1280 | 640 | 1280 | 1280 | 320 | 320 | 320 | 320 | 160 | 160 | YES |
| 8 | 75 | F | Bivalent | 2560 | 10240 | 226 | 2560 | 320 | 2560 | 113 | 640 | 57 | 640 | 80 | 320 | NO |
| 9 | 64 | M | Bivalent | 320 | 1810 | 40 | 640 | 57 | 905 | 20 | 226 | 10 | 320 | 10 | 57 | YES |
| 10 | 65 | F | Bivalent | 640 | 2560 | 40 | 160 | 80 | 320 | 20 | 80 | 28 | 80 | 10 | 40 | YES |
| 11 | 59 | M | Bivalent | 226 | 2560 | 10 | 320 | 20 | 226 | 20 | 160 | 10 | 80 | 10 | 20 | NO |
| 12 | 71 | F | Bivalent | 640 | 40960 | 160 | 20480 | 226 | 14482 | 20 | 1810 | 160 | 2560 | 10 | 640 | YES |
| 13 | 77 | M | Bivalent | 10240 | 10240 | 2560 | 3620 | 3620 | 5120 | 1280 | 1280 | 1810 | 2560 | 320 | 640 | YES |
| 14 | 68 | F | Bivalent | 640 | 2560 | 57 | 640 | 40 | 905 | 10 | 113 | 10 | 226 | 10 | 80 | NO |
| 15 | 60 | F | Bivalent | 1280 | 7241 | 320 | 1810 | 160 | 320 | 10 | 80 | 40 | 160 | 10 | 80 | YES |
| 16 | 61 | M | Bivalent | 80 | 1280 | 10 | 226 | 10 | 320 | 10 | 57 | 10 | 80 | 10 | 40 | NO |
| 17 | 69 | M | Bivalent | 1810 | 5120 | 80 | 3620 | 160 | 2560 | 40 | 640 | 10 | 453 | 20 | 320 | YES |
| 18 | 79 | M | Bivalent | 160 | 7241 | 10 | 2560 | 10 | 3620 | 10 | 1810 | 10 | 453 | 10 | 160 | NO |
| 19 | 57 | F | Bivalent | 320 | 2560 | 113 | 1280 | 80 | 2560 | 28 | 1280 | 10 | 640 | 10 | 320 | YES |
| 20 | 68 | M | Bivalent | 320 | 1280 | 57 | 320 | 113 | 640 | 40 | 160 | 40 | 226 | 10 | 40 | YES |
| 21 | 63 | M | Bivalent | 160 | 1280 | 40 | 320 | 113 | 453 | 20 | 113 | 20 | 160 | 20 | 113 | NO |
| 22 | 57 | M | Bivalent | 453 | 10240 | 28 | 3620 | 57 | 2560 | 28 | 640 | 10 | 320 | 10 | 160 | NO |
| 23 | 58 | M | Bivalent | 640 | 2560 | 40 | 1280 | 57 | 905 | 10 | 160 | 10 | 160 | 10 | 40 | NO |
| 24 | 66 | M | Bivalent | 226 | 20480 | 20 | 10240 | 40 | 10240 | 10 | 1280 | 10 | 2560 | 10 | 1280 | NO |
| 25 | 60 | F | Bivalent | 1280 | 5120 | 160 | 2560 | 453 | 2560 | 320 | 1280 | 80 | 640 | 40 | 320 | YES |
| 26 | 74 | M | Bivalent | 5120 | 10240 | 640 | 2560 | 1280 | 2560 | 57 | 320 | 1280 | 2560 | 28 | 80 | YES |
| 27 | 58 | M | Bivalent | 1280 | 5120 | 320 | 1280 | 320 | 640 | 80 | 160 | 80 | 320 | 28 | 57 | YES |
| 28 | 69 | M | Bivalent | 453 | 2560 | 20 | 640 | 40 | 640 | 10 | 80 | 10 | 160 | 10 | 40 | NO |
| 29 | 63 | M | Bivalent | 453 | 1810 | 10 | 160 | 10 | 226 | 10 | 160 | 10 | 57 | 10 | 20 | NO |
| 30 | 75 | F | Bivalent | 160 | 640 | 10 | 160 | 10 | 320 | 10 | 80 | 10 | 160 | 10 | 40 | NO |
| 31 | 66 | F | Bivalent | 1280 | 3620 | 226 | 640 | 320 | 905 | 160 | 160 | 80 | 160 | 80 | 113 | YES |
| 32 | 59 | F | Bivalent | 2560 | 3620 | 640 | 1280 | 905 | 2560 | 80 | 320 | 453 | 640 | 80 | 160 | YES |
| [@]33 | 63 | M | Bivalent | N/A | N/A | N/A | N/A | 10 | 453 | 10 | 80 | 10 | 160 | 10 | 40 | NO |
| 34 | 72 | M | Bivalent | 57 | 320 | 10 | 113 | 10 | 160 | 10 | 20 | 10 | 20 | 10 | 20 | NO |
| 35 | 58 | M | Bivalent | 2560 | 5120 | 453 | 905 | 320 | 1280 | 160 | 320 | 40 | 160 | 28 | 80 | YES |
| 36 | 73 | M | Bivalent | 20 | 320 | 10 | 113 | 10 | 40 | 10 | 10 | 10 | 10 | 10 | 10 | NO |
| 37 | 69 | F | Bivalent | 113 | 2560 | 10 | 640 | 10 | 320 | 10 | 40 | 10 | 226 | 10 | 40 | NO |
| 38 | 73 | F | Bivalent | 80 | 1280 | 10 | 80 | 10 | 160 | 10 | 28 | 10 | 57 | 10 | 20 | NO |
| 39 | 62 | M | Bivalent | 1810 | 1810 | 57 | 905 | 80 | 1280 | 20 | 160 | 10 | 160 | 20 | 80 | YES |
| 40 | N/A | N/A | Bivalent | N/A | N/A | N/A | N/A | N/A | N/A | N/A | N/A | N/A | N/A | N/A | N/A | NA |

*Individual FFRNT$_{50}$ value is the geometric mean of duplicate FFRNT results.
^FFRNT$_{50}$ of <20 was treated as 10 for plot purpose and statistical analysis.
[$]Not available (N/A)

Supplementary Table 3. $FFRNT_{50}$ values of all subjects received BNT162b2 monovalent booster.

| ID | AGE (years) | SEX | Booster | *$FFRNT_{50}$ | | | | | | | | | | | | With infection at baseline (pre Dose 4) |
|---|---|---|---|---|---|---|---|---|---|---|---|---|---|---|---|---|
| | | | | USA-WA1/2020 | | BA.4/5-spike | | BA.4.6-spike | | BA.2.75.2-spike | | BQ.1.1-spike | | XBB.1-spike | | |
| | | | | Pre | 1MPD4 | Pre | 1MPD4 | Pre | 1MPD4 | Pre | 1MPD4 | Pre | 1MPD4 | Pre | 1MPD4 | |
| 1 | 63 | M | BNT162b2 | 640 | 1810 | 57 | 320 | 113 | 320 | 40 | 160 | 40 | 160 | 20 | 80 | YES |
| 2 | 71 | M | BNT162b2 | 5120 | 7241 | 905 | 640 | 905 | 905 | 640 | 640 | 160 | 320 | 320 | 320 | YES |
| 3 | 67 | M | BNT162b2 | 10240 | 10240 | 1810 | 3620 | 2560 | 1810 | 1280 | 905 | 160 | 226 | 80 | 226 | YES |
| 4 | 64 | M | BNT162b2 | 7241 | 5120 | 640 | 640 | 905 | 1280 | 320 | 453 | 113 | 160 | 80 | 160 | YES |
| 5 | 68 | M | BNT162b2 | 5120 | 10240 | 226 | 640 | 320 | 640 | 160 | 453 | 80 | 160 | 40 | 113 | YES |
| 6 | 56 | M | BNT162b2 | 2560 | 7241 | 160 | 2560 | 640 | 2560 | 320 | 640 | 80 | 320 | 160 | 160 | YES |
| 7 | 66 | M | BNT162b2 | 3620 | 10240 | 160 | 1280 | 226 | 640 | 80 | 320 | 80 | 160 | 40 | 80 | YES |
| 8 | 57 | F | BNT162b2 | 453 | 640 | 40 | 40 | 28 | 40 | ^10 | 20 | 10 | 20 | 10 | 10 | NO |
| 9 | 61 | F | BNT162b2 | 2560 | 10240 | 320 | 1280 | 453 | 1280 | 160 | 320 | 113 | 160 | 80 | 226 | YES |
| 10 | 63 | F | BNT162b2 | 5120 | 7241 | 226 | 320 | 113 | 320 | 160 | 640 | 40 | 160 | 40 | 80 | YES |
| 11 | 67 | F | BNT162b2 | 10 | 1280 | 10 | 160 | 10 | 80 | 10 | 20 | 10 | 20 | 10 | **10** | NO |
| 12 | 72 | M | BNT162b2 | 20480 | 20480 | 2560 | 2560 | 1280 | 1810 | 453 | 640 | 160 | 160 | 160 | 226 | YES |
| 13 | 64 | F | BNT162b2 | 1280 | 1280 | 113 | 320 | 113 | 160 | 40 | 57 | **10** | 20 | 10 | 40 | NO |
| 14 | 56 | F | BNT162b2 | 10240 | 10240 | 1280 | 1810 | 640 | 2560 | 320 | 1280 | 160 | 640 | 160 | 640 | YES |
| 15 | 58 | M | BNT162b2 | 453 | 1280 | 20 | 80 | 28 | 80 | 10 | 40 | 10 | 20 | 10 | 20 | NO |
| 16 | 57 | F | BNT162b2 | 640 | 2560 | 113 | 160 | 226 | 320 | 80 | 160 | 40 | 80 | 57 | 80 | NO |
| 17 | 69 | M | BNT162b2 | 20 | 1280 | 10 | 113 | 10 | 57 | 10 | 40 | 10 | **10** | 10 | 10 | NO |
| 18 | 65 | F | BNT162b2 | 7241 | 5120 | 160 | 160 | 160 | 320 | 28 | 40 | 160 | 80 | 20 | 20 | NO |
| 19 | 67 | M | BNT162b2 | 640 | 1280 | 40 | 160 | 80 | 80 | 40 | 40 | 20 | 28 | 20 | 20 | YES |
| 20 | 68 | F | BNT162b2 | 226 | 1810 | 20 | 320 | 113 | 160 | 20 | 40 | **10** | 20 | 10 | 10 | NO |
| 21 | 70 | F | BNT162b2 | 5120 | 7241 | 226 | 453 | 320 | 453 | 113 | 160 | 80 | 160 | 80 | 80 | YES |
| 22 | 70 | M | BNT162b2 | 1280 | 2560 | 226 | 320 | 320 | 640 | 226 | 320 | 40 | 80 | 40 | 80 | YES |
| 23 | 65 | M | BNT162b2 | 80 | 640 | 10 | 10 | 10 | 10 | 10 | 10 | 10 | 10 | 10 | 10 | NO |
| 24 | 60 | F | BNT162b2 | 57 | 1280 | 10 | 80 | 10 | 80 | 10 | 20 | 10 | 20 | 10 | 10 | YES |
| 25 | 73 | M | BNT162b2 | 113 | 640 | **10** | 80 | 40 | 113 | 10 | 20 | 10 | 40 | 10 | 20 | NO |
| 26 | 70 | F | BNT162b2 | 160 | 1280 | 10 | 80 | 10 | 80 | 10 | 40 | 10 | 20 | 10 | **10** | NO |
| 27 | 73 | F | BNT162b2 | 320 | 640 | 10 | 28 | **10** | 57 | 10 | 40 | 10 | 10 | 10 | **10** | NO |
| 28 | 56 | F | BNT162b2 | 5120 | 2560 | 160 | 160 | 113 | 160 | 160 | 160 | 57 | 80 | 40 | 80 | NO |
| 29 | 65 | F | BNT162b2 | 160 | 1280 | 10 | 40 | 10 | 20 | 10 | 10 | 10 | 10 | 10 | 10 | NO |
| 30 | 75 | M | BNT162b2 | 905 | 1280 | 113 | 80 | 160 | 226 | 80 | 113 | 40 | 40 | 20 | 20 | NO |
| 31 | 61 | F | BNT162b2 | 160 | 640 | **10** | 80 | 10 | 80 | 10 | 20 | 10 | 20 | 10 | 10 | NO |
| 32 | 73 | F | BNT162b2 | 2560 | 5120 | 320 | 320 | 160 | 320 | 40 | 80 | 80 | 160 | 40 | 40 | NO |
| 33 | 64 | F | BNT162b2 | 10240 | 10240 | 1280 | 2560 | 1810 | 1810 | 905 | 1280 | 453 | 640 | 320 | 453 | YES |
| 34 | 68 | M | BNT162b2 | 40 | 226 | 10 | 10 | 10 | 40 | 10 | 10 | 10 | 10 | 10 | 10 | NO |
| 35 | 79 | F | BNT162b2 | 453 | 2560 | 40 | 226 | 80 | 160 | 20 | 80 | 20 | 40 | 10 | 20 | NO |
| 36 | 56 | F | BNT162b2 | 5120 | 5120 | 640 | 1280 | 1280 | 1280 | 320 | 640 | 160 | 320 | 160 | 226 | YES |
| 37 | 61 | F | BNT162b2 | 320 | 2560 | 40 | 113 | 40 | 80 | **10** | 40 | 10 | 20 | 10 | 20 | NO |
| 38 | 67 | F | BNT162b2 | 453 | 1280 | 20 | 113 | 40 | 160 | 10 | 40 | **10** | 40 | 10 | 20 | YES |
| 39 | 62 | F | BNT162b2 | 905 | 3620 | 113 | 453 | 80 | 160 | 20 | 80 | 10 | 40 | 20 | 40 | YES |
| 40 | 69 | M | BNT162b2 | 640 | 2560 | 40 | 160 | 40 | 160 | **10** | 40 | **10** | 28 | 10 | 20 | YES |

*Individual $FFRNT_{50}$ value is the geometric mean of duplicate FFRNT results.
^$FFRNT_{50}$ of <20 was treated as 10 for plot purpose and statistical analysis.

Supplementary Table 4. Geometric mean neutralizing titers (GMT) and geometric mean fold rise (GMFR) from before to 1 month post-dose 4 for bivalent BA.4/5 or BNT162b2 monovalent booster.

| Vaccine Group | Subset | | FFRNT$_{50}$ | | | | | | | | | | | |
|---|---|---|---|---|---|---|---|---|---|---|---|---|---|---|
| | | | USA-WA1/2020 | | BA.4/5-spike | | BA.4.6-spike | | BA.2.75.2-spike | | BQ.1.1-spike | | XBB.1-spike | |
| | | | Pre | 1MPD4 | Pre | 1MPD4 | Pre | 1MPD4 | Pre | 1MPD4 | Pre | 1MPD4 | Pre | 1MPD4 |
| BNT162b2 Bivalent BA.4/BA.5 30 µg | All | $n | 37 | 37 | 37 | 37 | 38 | 38 | 38 | 38 | 38 | 38 | 38 | 38 |
| | | #GMT (95% CI) | 572.0 (357.4, 915.3) | 3327.7 (2317.6, 4778.0) | 65.7 (38.8, 111.3) | 855.6 (552.2, 1325.8) | 81.5 (47.0, 141.4) | 905.1 (589.5, 1389.6) | 29.3 (19.2, 44.8) | 195.6 (124.6, 306.9) | 29.1 (17.8, 47.4) | 252.4 (164.7, 387.0) | 17.6 (13.1, 23.7) | 84.5 (57.5, 124.2) |
| | | †GMFR (95% CI) | | 5.8 (4.0, 8.5) | | 13.0 (8.0, 21.1) | | 11.1 (7.1, 17.3) | | 6.7 (4.4, 10.2) | | 8.7 (5.7, 13.3) | | 4.8 (3.3, 6.9) |
| | With Prior Infection | $n | 19 | 19 | 19 | 19 | 19 | 19 | 19 | 19 | 19 | 19 | 19 | 19 |
| | | #GMT (95% CI) | 1376.9 (854.0, 2219.9) | 4847.4 (3228.6, 7277.6) | 206.6 (115.9, 368.3) | 1376.9 (820.1, 2311.6) | 281.6 (159.5, 497.2) | 1564.4 (938.2, 2608.5) | 62.0 (32.5, 118.0) | 325.9 (183.0, 580.5) | 74.4 (34.7, 159.4) | 444.4 (259.4, 761.3) | 26.8 (16.2, 44.2) | 130.9 (80.0, 214.3) |
| | | †GMFR (95% CI) | | 3.5 (2.1, 6.0) | | 6.7 (3.5, 12.7) | | 5.6 (3.1, 9.8) | | 5.3 (2.8, 9.8) | | 6.0 (3.2, 11.2) | | 4.9 (2.8, 8.5) |
| | Without Prior Infection | $n | 18 | 18 | 18 | 18 | 19 | 19 | 19 | 19 | 19 | 19 | 19 | 19 |
| | | #GMT (95% CI) | 226.3 (125.7, 407.5) | 2237.2 (1238.2, 4042.2) | 19.6 (12.7, 30.2) | 517.8 (260.5, 1029.5) | 23.6 (14.1, 39.4) | 523.6 (277.3, 988.8) | 13.9 (10.1, 19.1) | 117.3 (60.6, 227.2) | 11.4 (9.3, 13.9) | 143.4 (78.7, 261.3) | 11.6 (9.1, 14.7) | 54.5 (31.0, 95.9) |
| | | †GMFR (95% CI) | | 9.9 (6.2, 15.7) | | 26.4 (14.4, 48.3) | | 22.2 (12.7, 38.8) | | 8.4 (4.6, 15.5) | | 12.6 (7.1, 22.5) | | 4.7 (2.8, 7.9) |
| BNT162b2 30 µg | All | $n | 40 | 40 | 40 | 40 | 40 | 40 | 40 | 40 | 40 | 40 | 40 | 40 |
| | | #GMT (95% CI) | 874.3 (479.7, 1593.3) | 2604.7 (1863.6, 3640.7) | 82.1 (47.6, 141.8) | 236.3 (148.6, 375.7) | 101.1 (58.9, 173.5) | 232.2 (149.3, 361.1) | 48.0 (29.1, 79.0) | 99.3 (62.4, 158.1) | 31.4 (21.4, 45.9) | 58.1 (39.2, 86.1) | 27.1 (18.9, 38.8) | 41.4 (27.5, 62.3) |
| | | †GMFR (95% CI) | | 3.0 (2.1, 4.3) | | 2.9 (2.1, 3.9) | | 2.3 (1.9, 2.8) | | 2.1 (1.7, 2.5) | | 1.8 (1.6, 2.2) | | 1.5 (1.3, 1.8) |
| | With Prior Infection | $n | 20 | 20 | 20 | 20 | 20 | 20 | 20 | 20 | 20 | 20 | 20 | 20 |
| | | #GMT (95% CI) | 2516.0 (1291.8, 4900.4) | 5120.0 (3465.6, 7564.2) | 226.3 (110.4, 463.8) | 629.0 (371.2, 1066.0) | 283.4 (142.6, 563.2) | 586.9 (346.9, 993.0) | 125.5 (62.1, 253.9) | 264.5 (146.3, 478.0) | 59.6 (35.0, 101.5) | 132.2 (82.5, 212.0) | 54.6 (32.5, 92.0) | 98.5 (58.0, 167.3) |
| | | †GMFR (95% CI) | | 2.0 (1.4, 2.9) | | 2.8 (1.9, 4.1) | | 2.1 (1.5, 2.8) | | 2.1 (1.6, 2.7) | | 2.2 (1.8, 2.7) | | 1.8 (1.5, 2.2) |
| | Without Prior Infection | $n | 20 | 20 | 20 | 20 | 20 | 20 | 20 | 20 | 20 | 20 | 20 | 20 |
| | | #GMT (95% CI) | 303.8 (137.9, 669.3) | 1325.1 (924.2, 1900.1) | 29.8 (16.9, 52.5) | 88.8 (55.3, 142.6) | 36.1 (20.4, 63.6) | 91.9 (59.8, 141.1) | 18.3 (12.1, 27.7) | 37.3 (25.1, 55.4) | 16.5 (11.0, 24.8) | 25.5 (17.4, 37.4) | 13.4 (10.3, 17.5) | 17.4 (12.6, 24.1) |
| | | †GMFR (95% CI) | | 4.4 (2.3, 8.2) | | 3.0 (1.8, 4.9) | | 2.5 (1.9, 3.5) | | 2.0 (1.6, 2.6) | | 1.5 (1.2, 1.9) | | 1.3 (1.1, 1.6) |

$Number of participants with valid and determinate assay results for the specific assay.

#Geometric mean neutralizing titers (GMT) and 2-sided 95% confidence interval (CI) were calculated by exponentiating the mean logarithm of the titers and the corresponding CIs (based on the Student t distribution). Assay results below the LLOQ were set to 0.5 × LLOQ.

†Geometric mean fold rise (GMFR) from before to 1 month post-dose 4 and 2-sided 95% CI were calculated by exponentiating the mean logarithm of fold rises and the corresponding CIs (based on the Student t distribution). Assay results below the LLOQ were set to 0.5 × LLOQ.
